# Accurate sample deconvolution of pooled snRNA-seq using sex-dependent gene expression patterns

**DOI:** 10.1101/2024.11.29.626066

**Authors:** Guy M. Twa, Robert A. Phillips, Nathaniel J. Robinson, Jeremy J. Day

## Abstract

Single-nucleus RNA sequencing (snRNA-seq) technology offers unprecedented resolution for studying cell type-specific gene expression patterns. However, snRNA-seq poses high costs and technical limitations, often requiring the pooling of independent biological samples and the loss of individual sample-level data. Deconvolution of sample identity using inherent features would enable the incorporation of pooled barcoding and sequencing protocols, thereby increasing data throughput and analytical sample size without requiring increases in experimental sample size and sequencing costs. In this study, we demonstrate a proof of concept that sex-dependent gene expression patterns can be leveraged for the deconvolution of pooled snRNA-seq data. Using previously published snRNA-seq data from the rat ventral tegmental area, we trained a range of machine learning models to classify cell sex using genes differentially expressed in cells from male and female rats. Models that used sex-dependent gene expression predicted cell sex with high accuracy (93-95%) and outperformed simple classification models using only sex chromosome gene expression (88-90%). The generalizability of these models to other brain regions was assessed using an additional published data set from the rat nucleus accumbens. Within this data set, model performance remained highly accurate in cell sex classification (90-92% accuracy) with no additional training. This work provides a model for future snRNA-seq studies to perform sample deconvolution using a two-sex pooled sample sequencing design and benchmarks the performance of various machine learning approaches to deconvolve sample identification from inherent sample features.

## INTRODUCTION

Single nucleus RNA sequencing (snRNA-seq) technologies enable the investigation of cell type-specific gene expression patterns, which are highly relevant to brain health and disease mechanisms [1–3]. Sample size is a key factor in snRNA-seq experimental design, as the number of experimental samples directly influences experimental costs and statistical power [4–6]. Additionally, droplet-based snRNA-seq technologies often require pooling tissue samples from multiple animals to achieve appropriate nuclei concentrations for maximum capture. Without the ability to deconvolute these samples, pooling decreases the effective sample size and causes individual-level data loss, thereby increasing the experimental costs required for sufficient statistical power with current gold-standard pseudobulking analysis approaches [7]. Additionally, the loss of individual-level data prevents the correlation of genetic contributions to molecular, physiological, or behavioral observations. Therefore, techniques to efficiently deconvolve pooled sample data are necessary to improve the power and cost-effectiveness of snRNA-seq technologies.

Currently, there are two popular methods for pooled sample deconvolution: nucleus hashing and genotype-based multiplexing [5,8,9]. Nuclei hashing requires tagging individual sample nuclei with oligonucleotide barcoded antibodies before pooling with other samples in sequence library preparation [9]. Following sequencing, cells are assigned to their respective samples based on the presence of detected hashing tags. This approach is limited by ambient hashing tag signal in suspension and attachment of hashing antibodies to lysis debris, both of which can make sample assignment non-trivial and noisy [9,10]. Genotype-based multiplexing deconvolutes cells by assigning them to samples based on shared genomic variants observed in sequenced nuclei libraries and sample genotypes [8]. This approach requires the collection of an additional data modality (sample genotype) and is limited by insufficient coverage of variant loci in snRNA-seq data. Multiplexing strategies utilizing both techniques may allow each to compensate for the other’s weaknesses [11,12]. However, this complicates sample processing by compounding sample preparation and data analysis requirements.

Here, we outline a model of pooled sample deconvolution centered on the classification of cell sex by sex-dependent transcriptome features inherent within the data, enabling sample assignment without requiring additional data modalities or sample preprocessing. We demonstrate that cell sex can be reliably recovered from snRNA-seq data by machine learning (ML) models, providing a proof of concept for sex-based sample pooling and deconvolution. This approach requires pooling nuclei of two animals of different sexes within a single microfluidic well and *post hoc* model-assisted cell sex classification for sample deconvolution. To determine the feasibility of ML models in cell sex classification, we trained and evaluated the performance of several widely used models with a range of complexities. When applied to previously published snRNA-seq data from the ventral tegmental area (VTA) of the rat brain, models trained using sex-dependent differentially expressed genes (DEGs) accurately classified up to 97% of cells. Models maintained high performance (>90% accuracy) when applied to independent snRNA-seq data from the rat nucleus accumbens (NAc), demonstrating their generalizability to distinct cell types in a different brain area. This analysis supports the pooling and deconvolution of samples in snRNA-seq assays based on sex, effectively halving experimental costs and doubling analytical power.

## RESULTS

### Determination of sex-dependent transcriptome features

To learn rules for predicting cell sex using transcriptome features, ML models require a relevant ground-truth training set of cells with known sex. To this end, we used previously published snRNA-seq data from the rat VTA containing 22,149 cells and 16 transcriptionally defined cell types (**Fig 1b**) [13]. This data set was obtained from pooled male or female samples assigned to separate 10X GEM wells and contained roughly equal numbers of male and female cells of each annotated cell type (**Fig 1c,d**). Cells were split into training and testing partitions of 70% and 30%, respectively (15,519:6,630), while maintaining cell type and cell sex ratios (**Fig. 1e,f**). The testing partition was set aside until model evaluation, and we leveraged the training partition to determine relevant sex-dependent transcriptome features and train classification models.

**Figure 1.**
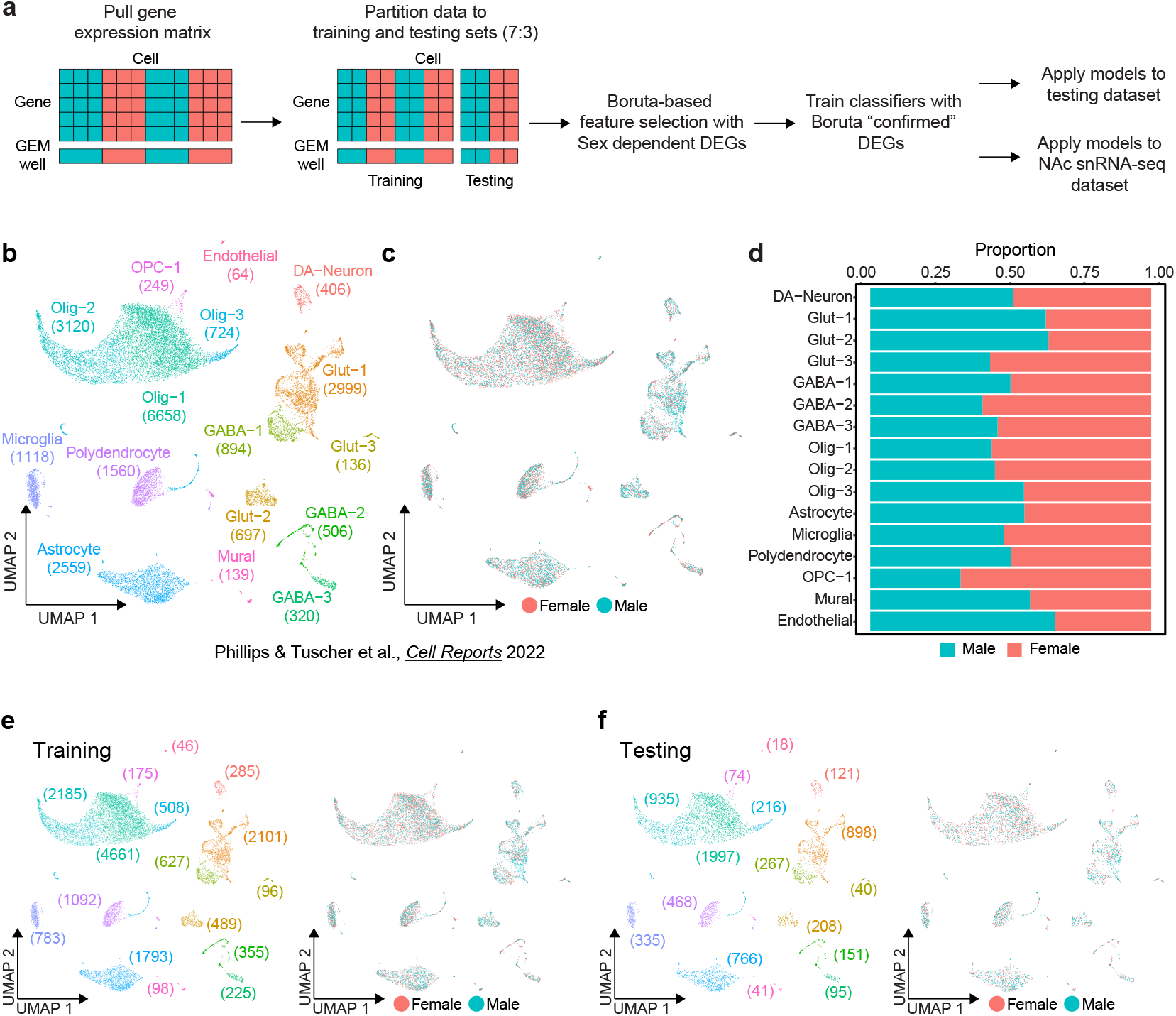
Sex classification model training workflow and ground truth data set. a, Schematic of classifier workflow including data preparation, feature selection, model training, and performance evaluation. b, UMAP of previously published rat ventral tegmental area (VTA) single nucleus RNA-seq data set including annotation of transcriptionally defined cell types (22,149 cells, 16 cell types) c, Distribution of sexes within UMAP. d, Stacked bar chart of the proportion of each sex within each cell type. e, UMAP of the training partition (70%) of the dataset with distributions of cell types (left) and sexes (right). f, UMAP of the testing partition (30%) of the VTA dataset with distributions of cell types (left) and sexes (right).

Selecting informative variables for class prediction is essential for efficient model training and high performance. To choose the transcriptome features most relevant for cell sex classification, we implemented two successive feature selection steps: sex-dependent differentially expressed gene (DEG) testing and Boruta feature selection (**Fig. 2a**) [14]. Of 23,733 detected genes, DEG testing identified 741 genes with significant sex-biased expression in at least one cell type (FDR < 0.05) (**Fig. 2b**). Sex-dependent directionality of expression of these genes was broadly replicated across cell types. To refine the set of genes for prediction models, we applied the Boruta algorithm, which identifies essential features by comparing their impact on classification accuracy to shuffled “shadow” features [14]. Boruta rejected 439 genes and confirmed 302 as more important than shadow features (**Fig 2c**). Genes with known sex-dependent gene expression, such as X chromosome genes *Xist* and *Kdm6a* and Y chromosome genes *Uty* (*ENSRNOG00000060617*) and *Ddx3*, replicated sex-specific expression patterns in our data and were some of the most important features selected by Boruta (**Fig 2c,d**). Additionally, the long non-coding RNA *ENSRNOG00000065796* adjacent to the *Xist* locus exhibited strong sex-dependent expression. Conversely, genes expressed sparsely without prior evidence of sex-biased transcription were rejected

**Figure 2.**
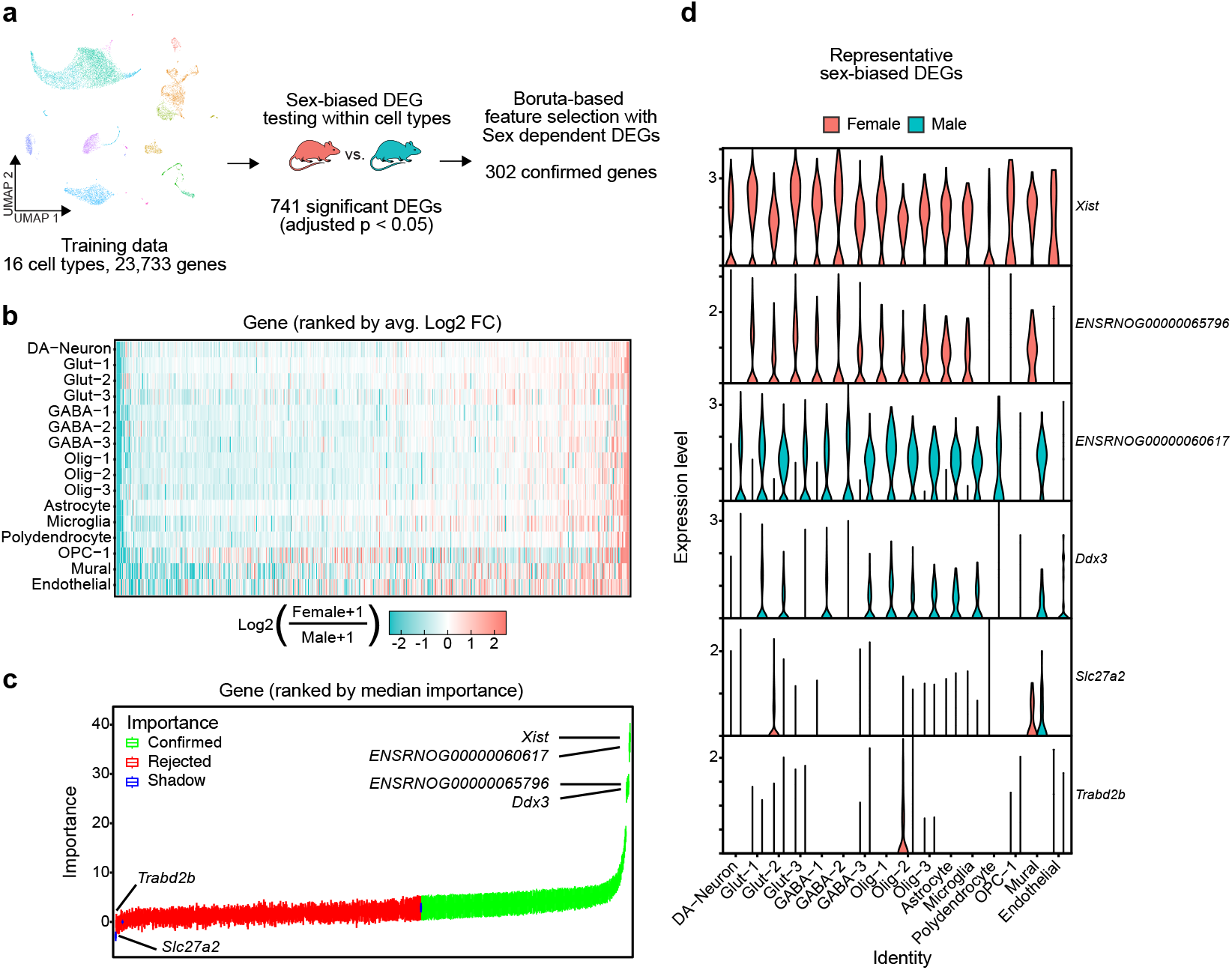
302 Sex-dependent DEGs confirmed as important features for cell sex classification model **a**, Schematic of feature selection using the VTA training data partition. **b**, Heatmap of log2 fold changes of genes significantly differentially expressed (adjusted p < 0.05) in at least one cell type (741 genes), color scale is capped from -2.5 to 2.5. Genes are ordered along the x-axis by ranked mean log2 fold change across cell types. **c**, Boxplots of gene importance scores as determined through 3000 iterations of the Boruta algorithm. Genes are ordered along the x-axis by median importance score. Boxplots are colored by the final importance decision by Boruta. (”Shadow” features are generated by Boruta from randomly shuffled gene data, used to evaluate the importance of real feature data). **d**, Violin plots of gene expression levels, within cell type split by sex, of the two highest confirmed importance female (*Xist, ENSRNOG00000065796*) and male (*ENSRNOG00000060617* (*Uty*), *Ddx3*) biased, and the two lowest importance rejected (*Slc27a2, Trabd2b*) sex-dependent DEGs determined by Boruta.

(**Fig. 2c,d**). Together, these processes narrowed the set of predictor variables to 302 transcripts (∼1% of all detected genes), with sex-dependent transcription demonstrating a more significant impact on cell sex classification accuracy than null shadow features.

### Machine learning models perform accurate cell sex classification

A wide range of approaches have been developed for classification tasks. To assess the potential for various models in cell sex classification, we surveyed four common ML models of varying complexity: logistic regression (LR), random forest (RF), support vector machine (SVM), and neural network multilayer-perceptron (MLP) models [15–22]. Additionally, we prepared two non-ML cell sex classification models based on the binary expression of sex chromosome genes: Xist, based on the expression of *Xist*, and Chr Y, based on the expression of chromosome Y genes. Together, these models represent various levels of complexity in approaches to cell sex classification strategies (**Fig. 3a**).

**Figure 3.**
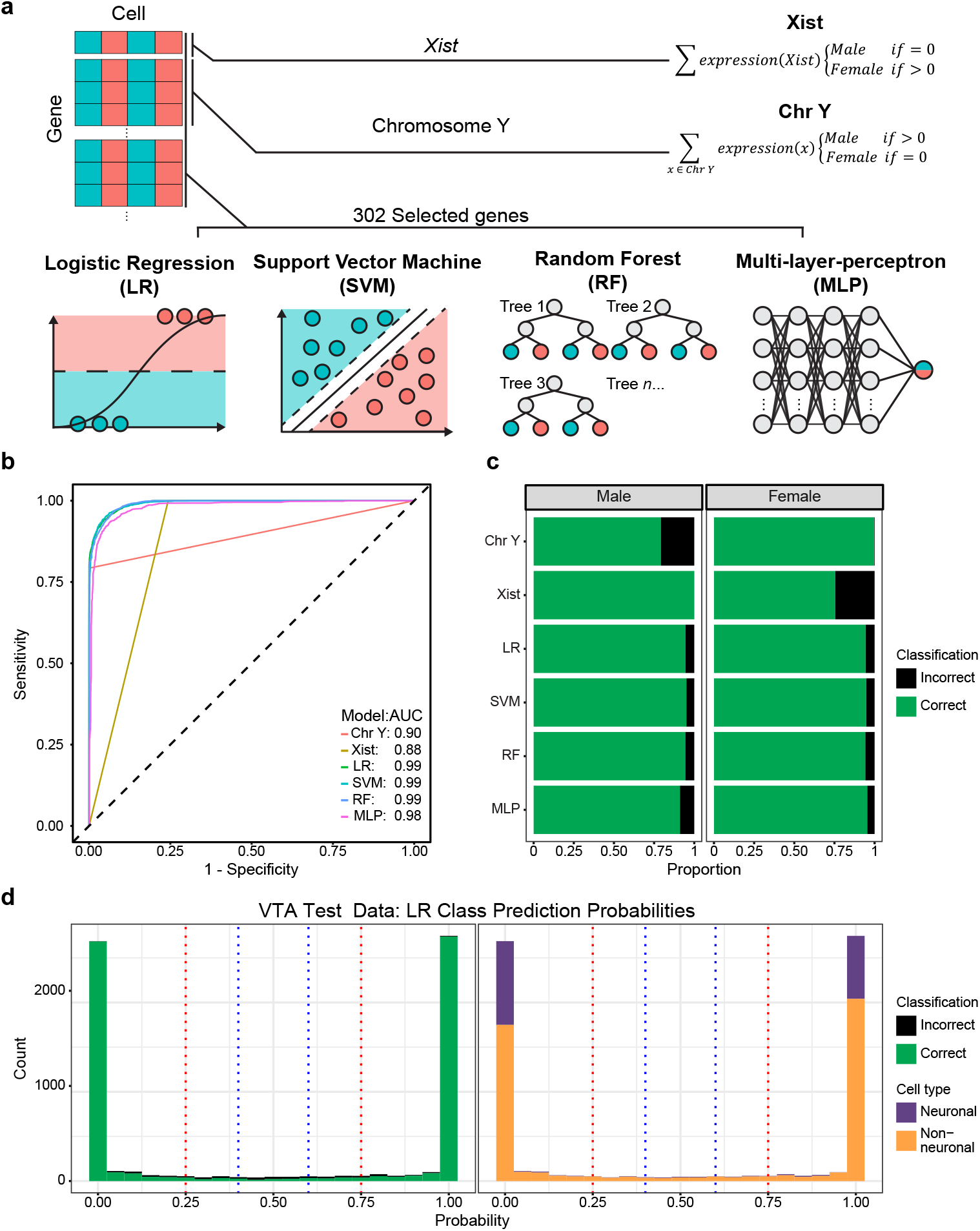
Machine learning models using selected genes outperform simple classifiers in sex classification. **a**, Schematic of models evaluated, the predic-tor genes used, and classification framework. **b**, Receiver operating characteristic curve of model sensitivity and specificity for classification of the VTA testing data partition. **c**, Stacked bar chart of the proportion of correct and incorrect classifications of the VTA testing data partition sexes. **d**, Histogram of logistic regression predicted cell sex probabilities (closer to 1 = high probability of being female, closer to 0 = high probability of being male) for the VTA test partition, bin size = 0.05. Blue and red dotted lines represent thresholds of 0.4-0.6 and 0.25-0.75 respectively. Bins are colored by the proportion of incorrect/correct classifications (left) and cell types (right).

When applied to the testing partition of the VTA data set, all ML models outperformed non-ML classification models based only on sex chromosome gene predictions. We evaluated models based on their overall accuracy, as well as sensitivity (true positive rate) and specificity (true negative rate) using the receiver operating characteristic curve (ROC). The area under the curve of the ROC (AUC-ROC) provides a quantitative measure of how well a classifier assigns observations, where a perfectly balanced classifier has an AUC-ROC of 1 and a random classifier has an AUC-ROC of 0.5. ML models achieved high overall accuracy (93-95%) and had balanced sensitivity and specificity (0.98-0.99 AUC-ROC) in cell sex classification (**Fig. 3b,c Table 1**). LR and SVM models outperformed all other methods (overall accuracies ∼95%, AUC-ROCs 0.99), despite the SVM model taking nearly 2,000x longer to train (**Fig. 3b,c, Table 1**). MLP and RF models only slightly underperformed LR and SVM models, with overall accuracies of 93% and 94%% and AUC-ROCs of 0.98 and 0.99, respectively. Non-ML classifiers suffered from comparatively poor accuracy, sensitivity, and specificity relative to the ML models. The Xist model was perfectly sensitive (misclassifying no male cells), but lacked specificity (misclassifying 25% of female cells; **Fig. 3b,c**). Conversely, the Chr Y model was nearly perfectly specific, misclassifying only 13 female cells (potentially due to misaligned transcripts) and poorly sensitive, misclassifying 21% of male cells (**Fig. 3b,c**). The high proportion of misclassifications by the non-ML models highlights a potential disadvantage of relying on a few predictors, as snRNA-seq data is prone to sparsity and gene drop-out. Accuracy and the number of UMIs (unique molecular identifiers) were significantly positively associated with one another (*p* < 0.05) for all models (**Fig. S1, Table S1**). All ML models required fewer UMIs to reach a 95% predicted probability of correct classification compared to non-ML classifiers. Together, these results demonstrate the ability of ML models, utilizing a robust set of sex-dependent genes, to accurately classify snRNA-seq cell sex.

**Table 1.** Ventral tegmental area model performance. Cell sex classification of the ventral tegmental area (VTA) testing partition was performed using two non-ML and four ML models. Model performance was assessed by accuracy and AUC-ROC. Model: Name of classification model; Predictors: set of predictor variables in model; Training time: total model training time in hours (hr), minutes (min), and seconds (s); Overall: performance measured across all cells; Neuronal: performance measured across only neuronal cell types; Non-neuronal: performance measured across only non-neuronal cell types; AUC-ROC: area under the curve of the receiver operating characteristic curve; Accuracy: proportion of correct classifications out of all classifications.

| Model | Predictors | Training time (hr:min:s) | Overall |  | Neuronal |  | Non-Neuronal |  |
| --- | --- | --- | --- | --- | --- | --- | --- | --- |
|  |  |  | AUC-ROC | Accuracy | AUC-ROC | Accuracy | AUC-ROC | Accuracy |
| Xist | Xist | N/A | 0.878 | 0.876 | 0.944 | 0.949 | 0.859 | 0.850 |
| Chr Y | Chromosome Y genes | N/A | 0.896 | 0.896 | 0.965 | 0.957 | 0.865 | 0.873 |
| Logistic Regression (LR) | 302 Selected features | 00:00:29 | 0.992 | 0.945 | 0.998 | 0.975 | 0.989 | 0.934 |
| Support Vector Machine (SVM) | 302 Selected features | 15:24:51 | 0.992 | 0.950 | 0.998 | 0.976 | 0.988 | 0.940 |
| Random Forest (RF) | 302 Selected features | 09:37:56 | 0.992 | 0.943 | 0.999 | 0.977 | 0.988 | 0.931 |
| Multi-Layer-Perceptron (MLP) | 302 Selected features | 07:56:12 | 0.980 | 0.933 | 0.995 | 0.963 | 0.974 | 0.922 |

Machine learning models predict cell sex probabilities from 0 (likely male) to 1 (likely female), enabling us to refine accuracy by omitting tenuous predictions near the classification threshold of 0.5. We assessed the number of cells omitted and improvements to accuracy for two thresholds centered around 0.5: narrow (0.4-0.6) and wide (0.25-0.75) (**Fig. 3d**). We found that the accuracy of cell sex predictions within the narrow threshold was 49% as compared to 96% accuracy outside, and within the wide threshold was 57% as compared to 97% outside (**Table 2**). The omission of cells within the narrow and wide thresholds would exclude 2.8% to 7.3% of cells from sex prediction, respectively (**Table 2**). Notably, cells within these low-accuracy thresholds tended to be non-neuronal cells (**Table 2**). The replication of this trend across all models highlights a broader challenge in cell sex classification – accurate classification of non-neuronal cells (**Table 1**).

**Table 2.**
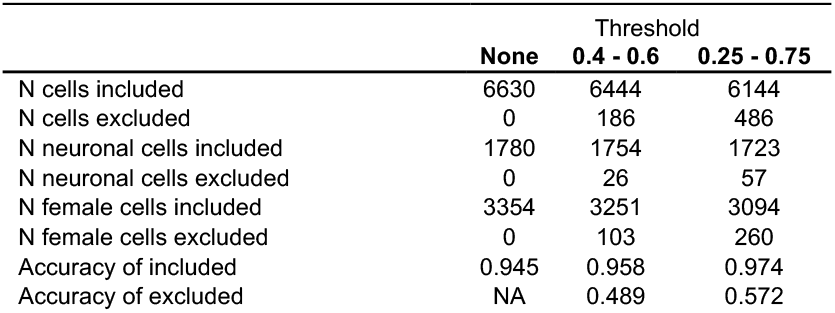
Predicted probability thresholding improves accuracy. Ventral tegmental area cells with tenuous (∼0.5) logistic regression (LR) predicted class probabilities were either retained or omitted using either narrow (0.4-0.6) or wide (0.25-0.75) thresholds. None: No omission by thresholding; 0.4-0.6: Omission of cells with class probabilities between 0.4 and 0.6; 0.25-0.75: Omission of cells with class probabilities between 0.25 and 0.75; N cells included: number of cells included after thresholding; N cells excluded: number of cells excluded after thresholding; N female cells included: number of female cells included after thresholding; N female cells included: number of female cells excluded after thresholding; Accuracy of included: Proportion of correct classifications out of all classifications made on included cells, outside of the threshold; Accuracy of excluded: Proportion of correct classifications out of all classifications made on excluded cells, inside of the threshold.

### Model performance is limited by information content in non-neuronal cell transcriptome

To explore the performance disparity between neuronal and non-neuronal cell populations, we assessed the performance of models explicitly trained for either cell class. Class-specific models were trained with the same workflow as pan-cellular models, using the VTA training dataset’s cell type-specific data partitions (**Fig. 4a**). Independent feature selection confirmed the importance of 59/129 neuronal and 244/656 non-neuronal DEGs, with 24 features overlapping. Despite differences in selected genes and training partitions, cell type-specific models performed largely similarly, although with slight decreases in performance, to pan-cellular models on neuronal and non-neuronal cell sex classification (**Fig. 4b, Table 3**). Losses in AUC-ROC and neuronal and non-neuronal classification accuracy by class-specific models were within 0.006 AUC-ROC and 1.7% overall accuracy (**Table 3)**. Only the neuronal MLP improved its performance, increasing 0.001 AUC-ROC and 0.4% accuracy over its pan-cellular counterpart. This result suggests that the disparity in cell class-specific model performance was not driven by an imbalance in the cell class composition of the training data but rather due to inherent limitations in the ability of gene expression in non-neuronal cells to be used for cell sex classification.

**Table 3.** Cell type-specific classification performance. Cell sex classification of ventral tegmental area (VTA) testing partition was performed using either neuronal or non-neuronal specific models. Cell type-specific predictor variable selection and model training was conducted using respective subsets of the VTA training partition. Model performance was assessed by accuracy and AUC-ROC. Model: Name of classification model; Predictors: the set of predictor variables in the model; Neuronal: performance of neuronal-specific models measured across neuronal cells only; Non-neuronal: performance of non-neuronal models measured across only non-neuronal cell types; AUC-ROC: area under the curve of the receiver operating characteristic curve; Accuracy: proportion of correct classifications out of all classifications.

| Model | Predictors | Neuronal |  | Non-Neuronal |  |
| --- | --- | --- | --- | --- | --- |
|  |  | AUC-ROC | Accuracy | AUC-ROC | Accuracy |
| Logistic Regression (LR) | 59 neuronal or 244 non-neuronal selected features | 0.998 | 0.974 | 0.985 | 0.926 |
| Support Vector Machine (SVM) | 59 neuronal or 244 non-neuronal selected features | 0.997 | 0.972 | 0.982 | 0.923 |
| Random Forest (RF) | 59 neuronal or 244 non-neuronal selected features | 0.998 | 0.976 | 0.984 | 0.917 |
| Multi-Layer-Perceptron (MLP) | 59 neuronal or 244 non-neuronal selected features | 0.997 | 0.967 | 0.973 | 0.914 |

**Figure 4.**
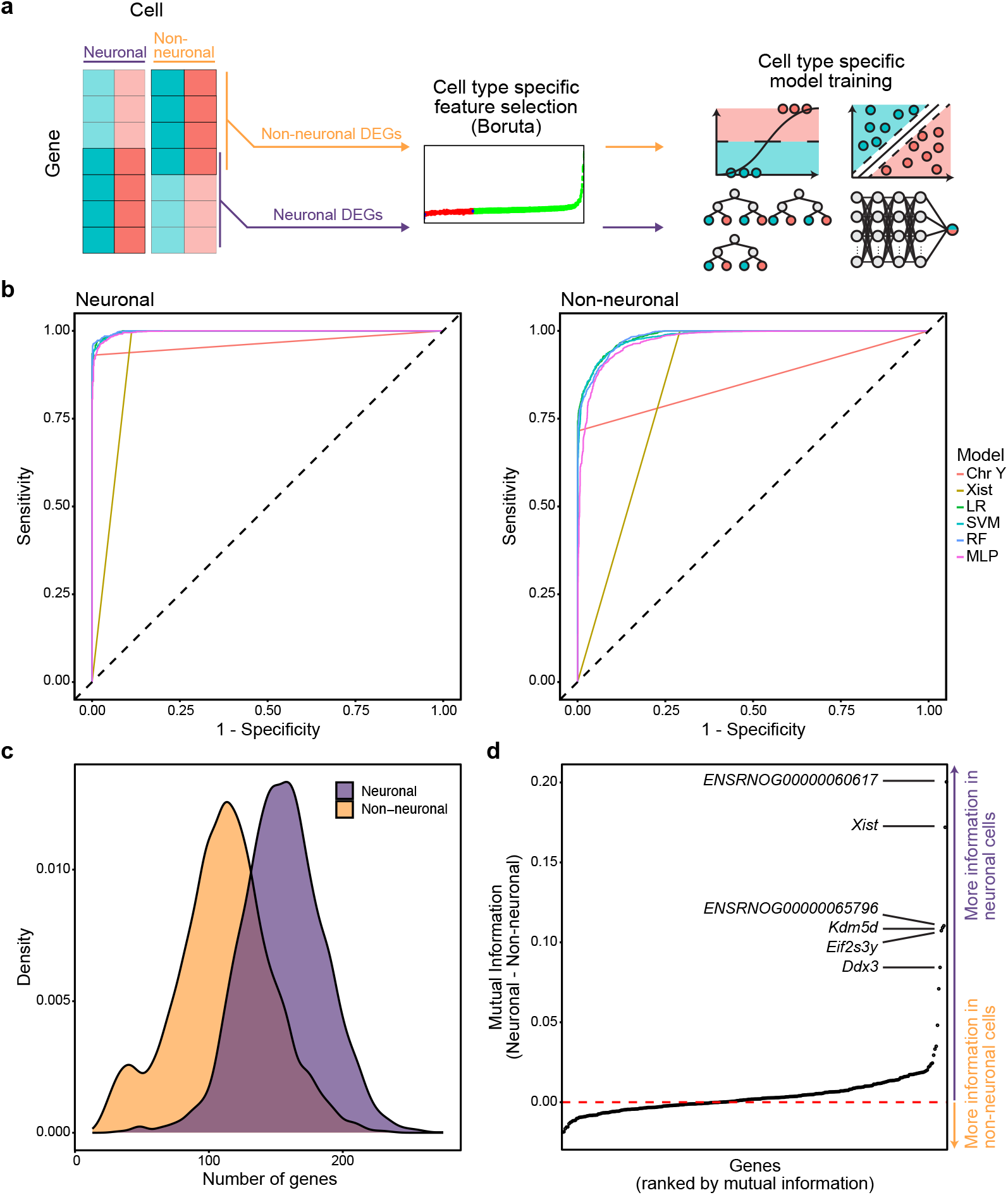
Cell type-specific models mirror the performance trends of models trained on all cells. **a**, Schema for cell type-specific model training strategy. **b**, Reciever operating characteristic curves of cell type-specific model sensitivity and specificity for classification of neuronal (left) and non-neuronal cells (right) of the VTA testing data partition. **c**, Distribution of the number of model predictor genes expressed in neuronal and non-neuronal cell populations of the VTA training data. **d**, Difference of mutual information score, calculated as mutual information of gene expression and sex, between neuronal and non-neuronal cells of the VTA training data partition.

We observed distinct neuronal and non-neuronal sex-dependent gene expression trends consistent with this hypothesis. In addition to having fewer detected genes on average, non-neuronal cells tend to express fewer model predictor genes than neuronal cells (**Fig. 4c**). Furthermore, we calculated the mutual information of the model gene’s expression and cell sex to quantify the information gained about cell sex by observing its expression in either neuronal or non-neuronal cell types [23,24]. Model predictor gene expression was more informative of cell sex in neuronal cells for 58% of genes (**Fig. 4d**). The six genes with the largest difference in mutual information (*ENSRNOG00000060617, Xist, ENSRNOG00000065796, Eif2s3y, Kdm5d, Ddx3*) were all located on sex chromosomes and conveyed more information about cell sex in neurons than non-neurons. Together, these observations underscore the inherent limitations in non-neuronal cell sex classification, as genes important for cell sex classification tend to be more sparsely detected in non-neuronal cells, and their expression tends to be less informative of cell sex.

### Model performance generalizes to independent data

Performance on new or unseen data, called generalizability, is a critical measure of a model’s utility. To validate that models learned robust cell sex classification rules that could be applied to another dataset, we measured the AUC-ROC and accuracy of model predictions in an orthogonal snRNA-seq experiment from the rat nucleus accumbens (NAc) [25]. Unlike the VTA, the NAc is composed mainly of GABAergic medium spiny neurons (in addition to cholinergic and GABAergic interneurons), yet also contains similar glial cell types [26]. This dataset includes 39,252 cells of 16 transcriptionally defined cell types from 32 rats (16M/16F) (**Fig. 5a,b**). While performance estimates were lower compared to the VTA dataset, all ML models achieved high overall accuracy (0.90-0.92) and balanced sensitivity and specificity (AUC-ROC 0.98-0.99) for NAc cell sex classification (**Fig. 5b,c, Table 4**). Of all ML models, the Random Forest model achieved the best overall performance with an AUC-ROC of 0.99 and an accuracy of 0.92 (**Fig. 5b,c, Table 4**). In comparison, the non-ML models remained poorly balanced for sensitivity and specificity, as shown by AUC-ROC scores of 0.92 for Xist and 0.88 for Chr Y classifiers (**Fig. 5b, Table 4**). However, the Xist model matched the ML models in terms of overall accuracy (0.91), even outperforming the MLP model (**Fig. 5b, Table 4**). Together, these results demonstrate that our feature selection and training strategies did not overfit models to the VTA data set, and that the resulting models are generalizable to cell sex classification of a distinct dataset from a different brain region.

**Table 4.** Nucleus accumbens classification performance. Cell sex classification of the nucleus accumbens dataset was performed using two non-ML and four ML models trained with the ventral tegmental area (VTA) training partition. Model performance was assessed by accuracy and AUC-ROC. Model: Name of classification model; Predictors: the set of predictor variables in the model; Overall: performance measured across all cells.

| Model | Predictors | Overall |  |
| --- | --- | --- | --- |
|  |  | AUC-ROC | Accuracy |
| Xist | Xist | 0.915 | 0.910 |
| Chr Y | Chromosome Y genes | 0.881 | 0.886 |
| Logistic Regression (LR) | 302 Selected features | 0.986 | 0.918 |
| Support Vector Machine (SVM) | 302 Selected features | 0.983 | 0.914 |
| Random Forest (RF) | 302 Selected features | 0.987 | 0.921 |
| Multi-Layer-Perceptron (MLP) | 302 Selected features | 0.975 | 0.903 |

**Figure 5.**
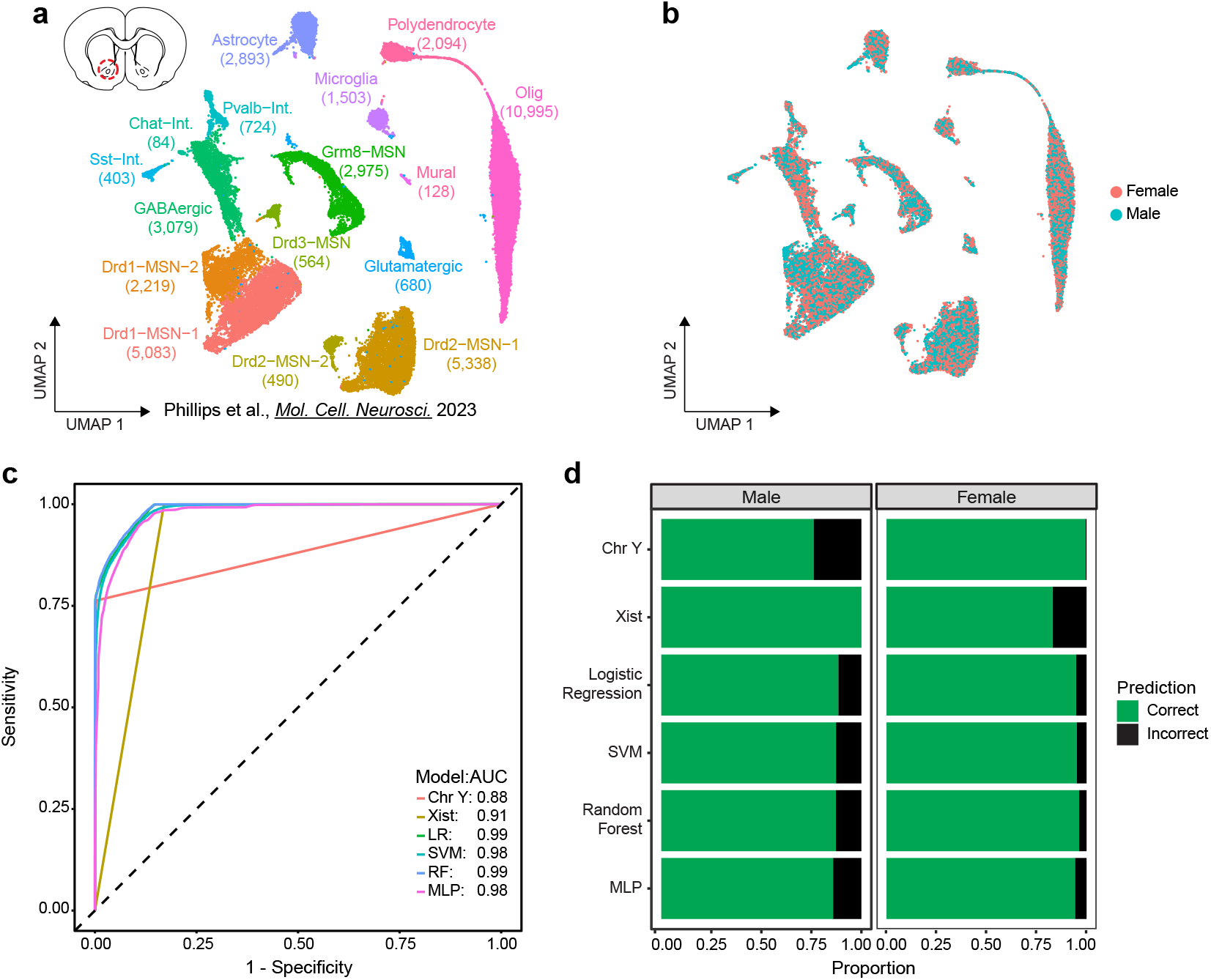
Sex classification models are generalizable to independent data sets. **a**, UMAP of previously published independent snRNA-seq data set of nucleus accumebns (NAc, top left diagram) samples from 32 (16 male, 16 female) rats including annotation of transcriptionally defined cell types (39,252 cells, 16 cell types). **b**, Distribution of sexes within UMAP. **c**, Reciever operating characteristic curve of VTA trained models’ sensitivity and specificity for classification of NAc cells. **d**, Stacked bar chart of proportion of correct and incorrect classifications of NAc sexes.

## DISCUSSION

Sample size and sequencing costs are practical limitations to snRNA-seq experimental design, especially when working with samples of limited starting material [4]. Pooling nuclei before sequencing is a popular method to overcome these limitations, but recovering individual-level data after sequencing poses additional challenges. Several methods for snRNA-seq demultiplexing have previously been proposed [5,8–10]. Our work expands the available toolkit for demultiplexing by providing a robust, low-cost method that leverages information inherent in the transcriptome to deconvolve pooled nuclei through sex prediction with machine learning models.

The use of sex-dependent transcripts is a key advantage of our demultiplexing strategy. In contrast, barcode hashing and genotype-based strategies require additional sample preparation steps before sequencing and involve non-trivial processing to assess sample identity post-sequencing [11,27]. Sex-based sample pooling eliminates the need for extra sample preparation or the collection of other modalities. Moreover, publicly available datasets can be used to identify sex-dependent transcriptional features and fit classification models (**Fig. 1, Fig. 2**). Highly accurate and lightweight machine learning models, such as logistic regression, can be quickly trained and applied to new data (**Fig. 3, Table 1**). Furthermore, we have demonstrated that these models generalize well to data from a distinct brain region, supporting their reusability across experiments (**Fig. 5**). Previous study designs with sex-balanced samples but pooled same-sex samples or different-sex samples that were not deconvolved might benefit from sex-dependent feature-based deconvolution with ML models [25,28]. Our sex-based classification approach thus complements and expands on previous strategies by leveraging orthogonal data inherent in sex-balanced snRNA-seq experiments.

Although this work directly evaluates the feasibility of sample deconvolution using sex-dependent transcriptome data, we also hypothesize that machine learning models could learn sex-dependent DNA accessibility features. This would enable their application in snATAC-seq experiments, expanding the potential for sex-based deconvolution in other modalities where sex differences in chromatin accessibility are observed. Additionally, this approach could be particularly valuable in multi-omic datasets, such as those combining snRNA-seq and snATAC-seq, where both modalities could be used synergistically to enhance sex prediction accuracy. However, further analysis and validation would be necessary to assess the robustness of such models with snATAC-seq data. Future work in this direction would represent a meaningful extension of sex-based deconvolution to additional sequencing modalities and multi-omic datasets.

Our evaluation of cell sex classification models identified poorer performance in non-neuronal cells as a potential limitation. While model performance remained above 87% accuracy, it was consistently lower than the performance observed with neuronal cells (**Fig. 4, Table 1, Table 2**). This discrepancy may be attributed to the observation that non-neuronal cells express fewer sex-predictive features and that expressed sex-predictive features tended to convey less information than in neuronal cell populations (**Fig. 4**). Additionally, our analysis revealed a significant relationship between UMI count and accuracy across all models (**Fig. S1**), suggesting that datasets with lower UMI counts – a common feature of non-neuronal populations in other brain snRNA-seq studies – may also exhibit reduced performance in sex-based classification [25,29,30].

One potential strategy to address this limitation is the integration of sex-based pooling and deconvolution with another demultiplexing strategy, such as barcode hashing, which does not exhibit the same cell-type constraints [9]. Using multiple orthogonal methods for sample deconvolution offers a practical way to resolve ambiguous sample assignments and identify potential doublets [5]. Integration of additional deconvolution approaches would permit future studies to take advantage of the complementary strengths of each method and mitigate their limitations [11,12].

In conclusion, we demonstrate that sex-dependent transcripts can be leveraged to train accurate cell sex prediction models, supporting the feasibility of sex-based pooling and sample demultiplexing. This work represents a meaningful expansion of previously proposed strategies for sample deconvolution. Our approach does not require additional sample preprocessing or modalities, and demultiplexing can be achieved using lightweight machine-learning models. Moreover, because it does not rely on additional data modalities, our method is highly compatible with other sample deconvolution strategies, allowing them to complement one another’s strengths. Accurate and easy-to-implement sex-based sample deconvolution enables future work to carry out less expensive and more powerful snRNA-seq analyses.

## METHODS

### Computational resources and environment

To maintain a consistent model training and evaluation environment, all analyses were run on UAB’s Cheaha research computing cluster (8 CPUs with 16 GB of RAM per CPU). All analyses used R version 4.2.0 for the x86_64-pc-linux-gnu platform, with a consistent random seed (set_seed(1234)).

### snRNA-seq data for model training and evaluation

To train and evaluate models, raw fastq files obtained from previous publications of rat snRNA-seq data were aligned to the Ensembl mRatBn7.2 (Rn7) genome with a custom version of the corresponding Ensembl annotation file (v105) using CellRanger (v6.1.2) [13, 25]. The custom annotation file was created to include the genome annotation data for *Xist*, absent in the Ensembl annotation file, from the NCBI RefSeq annotation file (GCF_015227675.2). Seurat (version 5.2.01) objects for each dataset were created for analyses, maintaining cell type annotations and dimensionality reductions from their original publications [31]. The genetic sex of nuclei was determined by the originating GEM (gel bead in emulsion) well used for nuclei capture, as sample nuclei of same-sex samples were pooled before capture. Analyses were conducted using RNA counts log-normalized and scaled by a factor of 10,000. Training and testing partitions of the VTA data set were created with a ratio of 7:3 by randomly assigning cells to either partition based on cell type and sex, maintaining distributions between partitions.

### Differential expression

Differentially expressed genes (DEGs) between male and female cells were determined for each cell type by the Wilcoxon Rank Sum test implemented in the FindMarkers() function from Seurat with default parameters. Significant DEGs were identified as those with Bonferroni-adjusted p-values < 0.05 in at least one cell type.

### Boruta feature selection

The Boruta feature selection algorithm further refined the set of significant DEGs by identifying genes with significant importance for cell sex classification within the VTA training partition. Importance was determined by iteratively comparing the importance, measured as Z-scores of the mean decrease accuracy measure, of real genes to a set of null “shadow features” constructed from shuffled gene counts using the Boruta() function from the Boruta package (version 8.0.0). At each iteration, genes with significantly lower scores are rejected and removed, while genes with significantly higher scores are confirmed and retained. After 3000 iterations, final decisions of genes with tentative importance were made using the TentativeRoughFix() Boruta function. The final set of genes with confirmed importance was used as predictive variables for cell sex classification models.

### Mutual Information

To compare the information shared between a gene’s expression and cell sex in neuronal and non-neuronal cell types, the mutual information of the two variables was calculated separately for the two broad cell types. Mutual information of transcript count data and cell sex for all selected model genes was calculated using the mmi.pw() function from the mpmi package in R (version 0.43.2.1) [24]. For calculation of gene’s expression and cell sex mutual information content in neuronal or non-neuronal populations, the VTA training partition was separated into neuronal (Glut-Neuron-1, Glut-Neuron-2, Glut-Neuron-3, GABA-Neuron-1, GABA-Neuron-2, GABA-Neuron-3, DA-Neuron) and non-neuronal (Olig-1, Olig-2, Olig-3, Astrocyte, Polydendrocyte, Microglia, OPC-Olig-1, Mural, Endothelial) populations.

### Model Training

To train cell sex prediction models, models were fit using the VTA training partition log-normalized gene counts as predictor variables and cell sex as the outcome variable. The Xist model made binary predictions of female or male cell sex based on the presence or absence of *Xist* counts. Reciprocally, the Chromosome Y classifier made binary predictions of male or female cell sex based on the presence or absence of counts from any gene on the Y chromosome. The logistic regression model was fit to the training data with the glm() function from the stats package (version 4.2.0). To assess training for a range of hyperparameters, the random forest, support vector machine, and multilayer perceptron models were trained with the train() function of the caret package (version 6.0-94) [32]. Training parameters were set to output cell sex class probabilities and perform 3x repeated 10-fold cross-validation using trainControl() with parameters method = “repeatedcv”, number = 10, repeats = 3, classProbs = T, allowParallel = T. Ranges for model hyperparameters for each model were specified using tuning grids defined for each model, described in detail below. Model accuracy was used to select the optimal configuration for each model.

#### Random forest

The random forest model was trained with method = “rf ” and a constant number of 1000 trees ntrees = 1000 as train() function parameters. To tune the number of variables randomly sampled at each split, the custom tuning grid assessed a range of “mtry” values from 2 to 334 by steps of 12. The final optimal model selected for accuracy used ntree = 1000, and mtry = 74.

#### Support vector machine

The support vector machine (SVM) model was trained with method = “svmRadial” as a parameter of the train() function to specify an SVM model with a radial basis function. A custom tuning grid of hyperparameters specifying “sigma” values from 0.0001 to 0.1, where steps increase by a factor of 10 with each subsequent term, and C values from 0.1 to 10 (0.1,0.2,0.5,1,1.5,2,5,10), was utilized for model training. The final optimal model, selected for its accuracy, used sigma = 1e-3 and C = 5.

#### Multilayer perceptron

A multi-layer perceptron with weight decay was trained using method = “mlpWeightDecayML” as train() function parameters. Model hyperparameters for training were tuned using a custom tuning grid specifying ranges for the number of nodes for each layer 1 (1,5,10,15,20), layer 2 (0,2,5,8,10), layer 3 (0,1,2,4,5), and weight decay `decay` values (0,0.05,0.1,0.15,0.2). Only hyperparameter configurations with decreasing nodes in successive layers were evaluated. The final optimal model, selected for its accuracy, was trained using the following parameters: layer1 = 20, layer2 = 10, layer3 = 4, and decay = 0.

#### Neuronal/Non-neuronal models

Before fitting cell type-specific models, the VTA training data partition was split into subsets for either neuronal (Glut-Neuron-1, Glut-Neuron-2, Glut-Neuron-3, GABA-Neuron-1, GABA-Neuron-2, GABA-Neuron-3, DA-Neuron) or non-neuronal (Olig-1, Olig-2, Olig-3, Astrocyte, Polydendrocyte, Microglia, OPC-Olig-1, Mural, Endothelial) cell type groups. Feature selection using Boruta was repeated for the significant DEGs within each cell type subset. LR, RF, SVM, and MLP models were fit as described above for neuronal and non-neuronal cell type subsets.

### Model evaluation

Model performance was evaluated using the VTA testing partition and the NAc dataset. Neuronal and non-neuronal models were assessed using respective VTA testing data partition subsets. The overall model classification accuracy was calculated as the number of correct classifications divided by the total number of classifications made. For all models, the receiver operating characteristic (ROC) curve and the area under the ROC (AUC-ROC) were used to evaluate the trade-off of sensitivity and specificity rates. ROC curve and AUC values were calculated using the roc() function of the pROC package in R. True-positive and false-positive rates were calculated using males as the “positive” class.

## DATA AVAILABILITY

All relevant data that support the findings of this study are available by request from the corresponding author (J.J.D.). Sequencing data that support the findings of this study are available in Gene Expression Omnibus. Accession numbers of specific datasets are outlined below. Ventral tegmental area snRNA-seq VTA: GSE168156 Nucleus accumbens snRNA-seq: GSE137763, GSE222418

Custom code can be found at https://github.com/Jeremy-Day-Lab/Twa_etal_2024

## ACKNOWLEDGEMENTS

We thank all current and former Day Lab members for their assistance and support. This work was supported by NIH grants DP1DA039650, R01MH114990, R01DA053743, and R01DA054714 and the McKnight Foundation Neurobiology of Brain Disorders Award (JJD).

## AUTHOR CONTRIBUTIONS

Conceptualization: RAP, JJD

Methodology: RAP, GT, NJR

Data Curation: GT, RAP

Formal Analysis: GT

Visualization: GT

Supervision: JJD

Funding acquisition: JJD

Writing – original draft: GT

Writing – review & editing: GT, RAP, NJR, JJD

---

**Figure S1.**
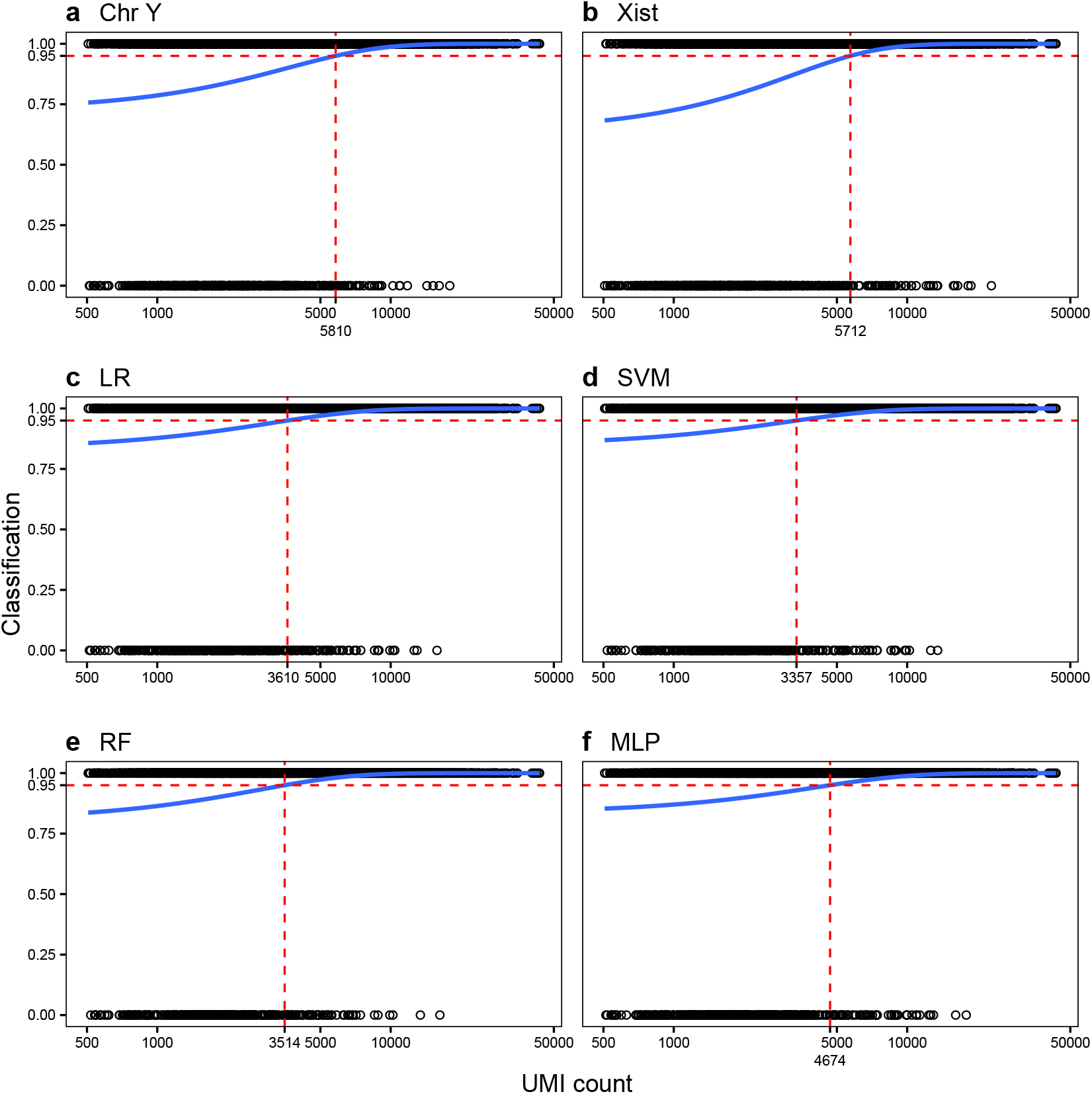
Increased RNA count per cell increases probability of correct classification for all models. Logistic regression of ventral tegmental area test partition cell UMI (unique molecular index) count and model classification (correct: 1, incorrect: 0) for chromosome Y (**a**), Xist (**b**), logistic regression (**c**), support vector machine (**d**), random forest (**e**), and multilayer-perceptron (**f**) models. Dashed red lines indicate the RNA count corresponding to a 95% probability of correct classification by a model.

**Table S1.** Increased UMI count improves the likelihood of accurate cell sex classification. The relationship between transcript UMI (unique molecular index) count and accuracy of model classification (incorrect: 0, correct: 1) in the ventral tegmental area test partition data was modeled using logistic regression for all cell sex classification models. Model: model classifications used to fit logistic regression; Intercept: fit logistic regression model intercept; UMI: fit logistic regression model coefficient for UMI count; Estimate: model estimate for either intercept or UMI terms; StdErr: standard error of either Intercept or UMI terms; Z-score: Estimate / Std Error; pval: p-value associated with the value Z-score column; 95% Probability Intercept: UMI value of where probability of correct classification reaches 95% as predicted by fit model.

| Model | Intercept |  |  |  | UMI count |  |  |  | 95% Probability Intercept |
| --- | --- | --- | --- | --- | --- | --- | --- | --- | --- |
|  | Estimate | StdErr | Z-score | pval | Estimate | StdErr | Z-score | pval |  |
| Chr Y | 0.963 | 0.076 | 12.682 | 7.44E-37 | 3.41E-04 | 2.38E-05 | 14.350 | 1.07E-46 | 5810 |
| Xist | 0.556 | 0.074 | 7.480 | 7.41E-14 | 4.18E-04 | 2.49E-05 | 16.821 | 1.72E-63 | 5712 |
| LR | 1.603 | 0.102 | 15.659 | 2.86E-55 | 3.72E-04 | 3.44E-05 | 10.814 | 2.97E-27 | 3610 |
| SVM | 1.706 | 0.106 | 16.025 | 8.51E-58 | 3.69E-04 | 3.58E-05 | 10.321 | 5.66E-25 | 3357 |
| RF | 1.413 | 0.104 | 13.529 | 1.05E-41 | 4.36E-04 | 3.74E-05 | 11.642 | 2.52E-31 | 3514 |
| MLP | 1.617 | 0.089 | 18.230 | 2.97E-74 | 2.84E-04 | 2.65E-05 | 10.734 | 7.04E-27 | 4674 |

